# A need for time-varying models to suppress artefacts of tACS in the M/EEG

**DOI:** 10.1101/2021.06.14.448446

**Authors:** Nicholas S. Bland

## Abstract

Rhythmic modulation of brain activity by transcranial alternating current stimulation (tACS) can entrain neural oscillations in a frequency- and phase-specific manner. However, large stimulation artefacts contaminate concurrent ‘online’ neuroimaging measures, including magneto- and electro-encephalography (M/EEG)—restricting most analyses to periods free from stimulation (‘offline’ aftereffects). While many published methods exist for removing artefacts of tACS from M/EEG recordings, they universally assume linear artefacts: either time-invariance (i.e., an artefact is a scaled version of itself from cycle to cycle) or sensor-invariance (i.e., artefacts are scaled versions of one another from sensor to sensor). However, heartbeat and respiration both nonlinearly modulate the amplitude and phase of these artefacts, predominantly via changes in scalp impedance. The spectral symmetry this introduces to the M/EEG spectra may lead to false-positive evidence for entrainment around the frequency of tACS, if not adequately suppressed. Good electrophysiological evidence for entrainment therefore requires that tACS artefacts are fully accounted for before comparing online spectra to a control (e.g., as might be observed during sham stimulation). Here I outline an approach to linearly solve templates for tACS artefacts, and demonstrate how event-locked perturbations to amplitude and phase can be introduced from simultaneous recordings of heartbeat and respiration—effectively forming time-varying models of tACS artefacts. These models are constructed for individual sensors, and can therefore be used in contexts with few EEG sensors and with no assumption of artefact collinearity. I also discuss the feasibility of this approach in the absence of simultaneous recordings of heartbeat and respiration traces.

## INTRODUCTION

The ability to modulate neural processes with non-invasive brain stimulation has furthered our understanding of brain–behaviour relationships in both health and disease (Polanía et al., 2018). Pairing techniques of non-invasive brain stimulation with neuroimaging has great scientific value (Bergmann et al., 2016; Soekadar et al., 2016), but the recovery of this information during stimulation (“online”) is challenging (Bland and Sale, 2019; Kasten and Herrmann, 2019). Techniques of *rhythmic* stimulation can entrain neural oscillations in a frequency- and phase-specific manner (Veniero et al., 2015; Fiene et al., 2020; Vieira et al., 2020; Huang et al., 2021), but they also contaminate the magneto- and electro-encephalogram (M/EEG) with large artefacts that complicate spectral analyses. For example, artefacts of transcranial alternating current stimulation (tACS; Herrmann et al., 2013)—where a low-intensity AC is applied between two or more scalp electrodes—are several orders of magnitude larger than the signals generated by endogenous neural activity in the EEG (**Figure 1**). This makes it difficult to directly observe neural entrainment, since the endogenous oscillations one typically wishes to recover occur at (or around) the frequency of tACS. Without fully accounting for these artefacts, spectral analyses are often restricted to recordings without active stimulation (offline; e.g., Zaehle et al., 2010), but this means that any neural entrainment must survive the period of stimulation in order to be detected (e.g., probing echoes of entrainment; see Geffen et al., 2021). Alternatively, EEG-correlates can be established and targeted across entirely separate experiments (e.g., Bland et al., 2018, 2020).

**Figure 1.**
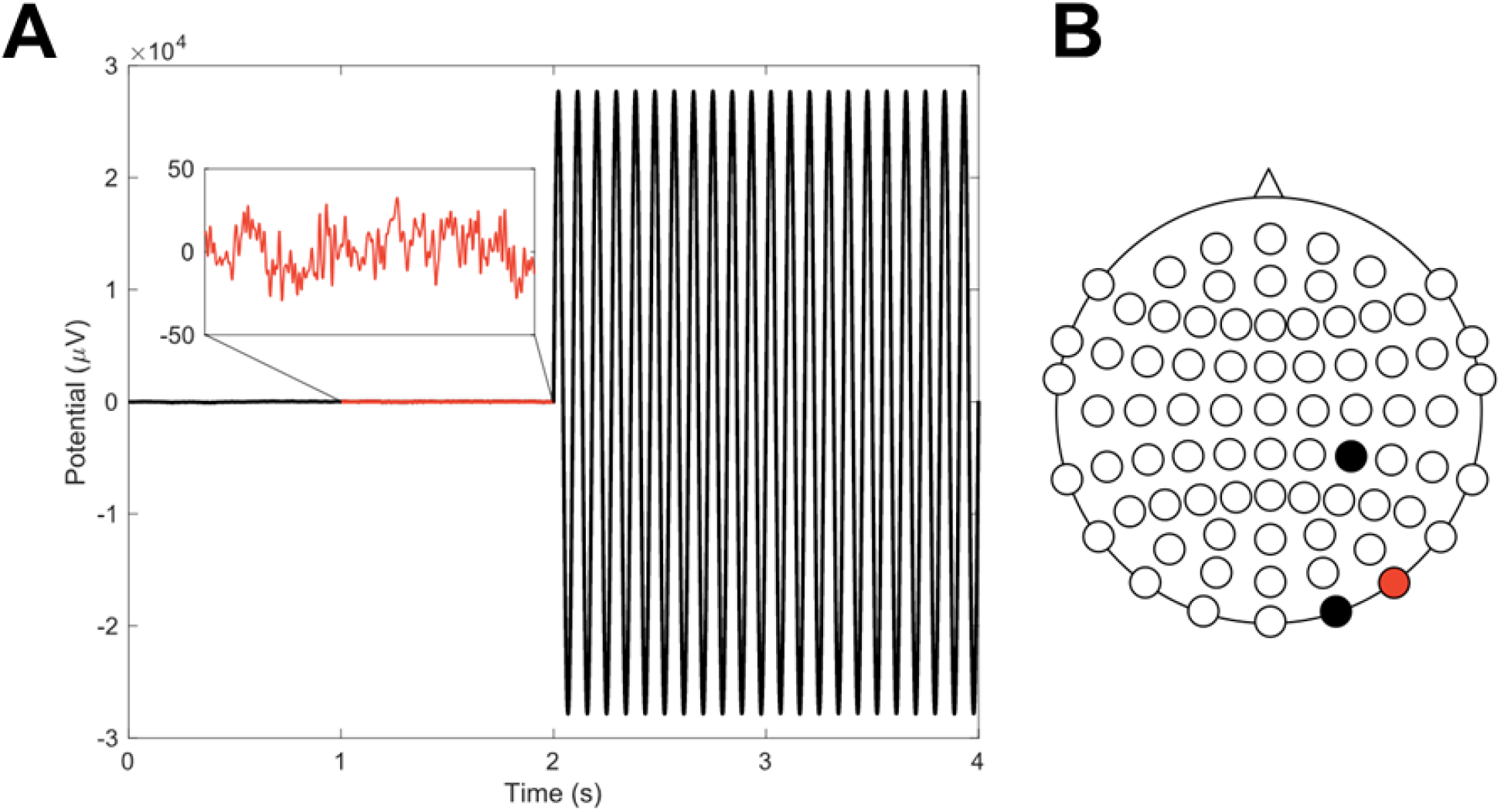
Artefacts of tACS contaminate electrophysiological recordings. **(A)** This EEG sensor contains an artefact of 11 Hz tACS several orders of magnitude larger than the endogenous oscillations (inset, red). A template of the artefact has been subtracted from the first half of the recording only. **(B)** Stimulation was applied at 1 mA peak-to-peak (black electrodes), with electrode PO10 (red) displayed.

Many methods exist for recovering the M/EEG from artefacts of tACS (Herrmann and Strüber, 2017; Kasten and Herrmann, 2019), with varying degrees of complexity and success. These methods include spatial filtering (beamforming, Neuling et al., 2015; Noury et al., 2016; Witkowski et al., 2016; ‘SASS,’ Haslacher et al., 2021), temporal filtering (notch, Voss et al., 2014; Hampel, Santos Monteiro et al., 2015), variations of template subtraction (Voss et al., 2014; Kohli and Casson, 2015, 2019; Noury et al., 2016; Dowsett and Taylor, 2017; Guggenberger and Gharabaghi, 2018), principal component analysis (PCA, Kohli and Casson, 2015; Santos Monteiro et al., 2015), and a combination of the latter two (Helfrich et al., 2014; Noury et al., 2016). Universally, these approaches assume *linear* stimulation artefacts— either time-invariance (i.e., an artefact is a scaled version of itself from cycle to cycle) or sensor-invariance (i.e., artefacts are scaled versions of one another from sensor to sensor). In recent years, the journal *NeuroImage* has been host to much commentary on the characterisation of tACS artefacts (e.g., Noury et al., 2016; Neuling et al., 2017; Noury and Siegel, 2017, 2018). It has been demonstrated that physiological rhythms in scalp impedance and head position (e.g., from heartbeat and respiration) nonlinearly modulate the amplitude (Noury et al., 2016) and phase (Noury and Siegel, 2017) of tACS artefacts— contaminating the power spectrum symmetrically around the frequency of stimulation (**Figure 2**). Since any residual tACS artefacts result in spectral symmetry close to where entrainment is presumed to occur, failure to fully account for this may result in false-positive evidence for neural entrainment (Noury and Siegel, 2018).

**Figure 2.**
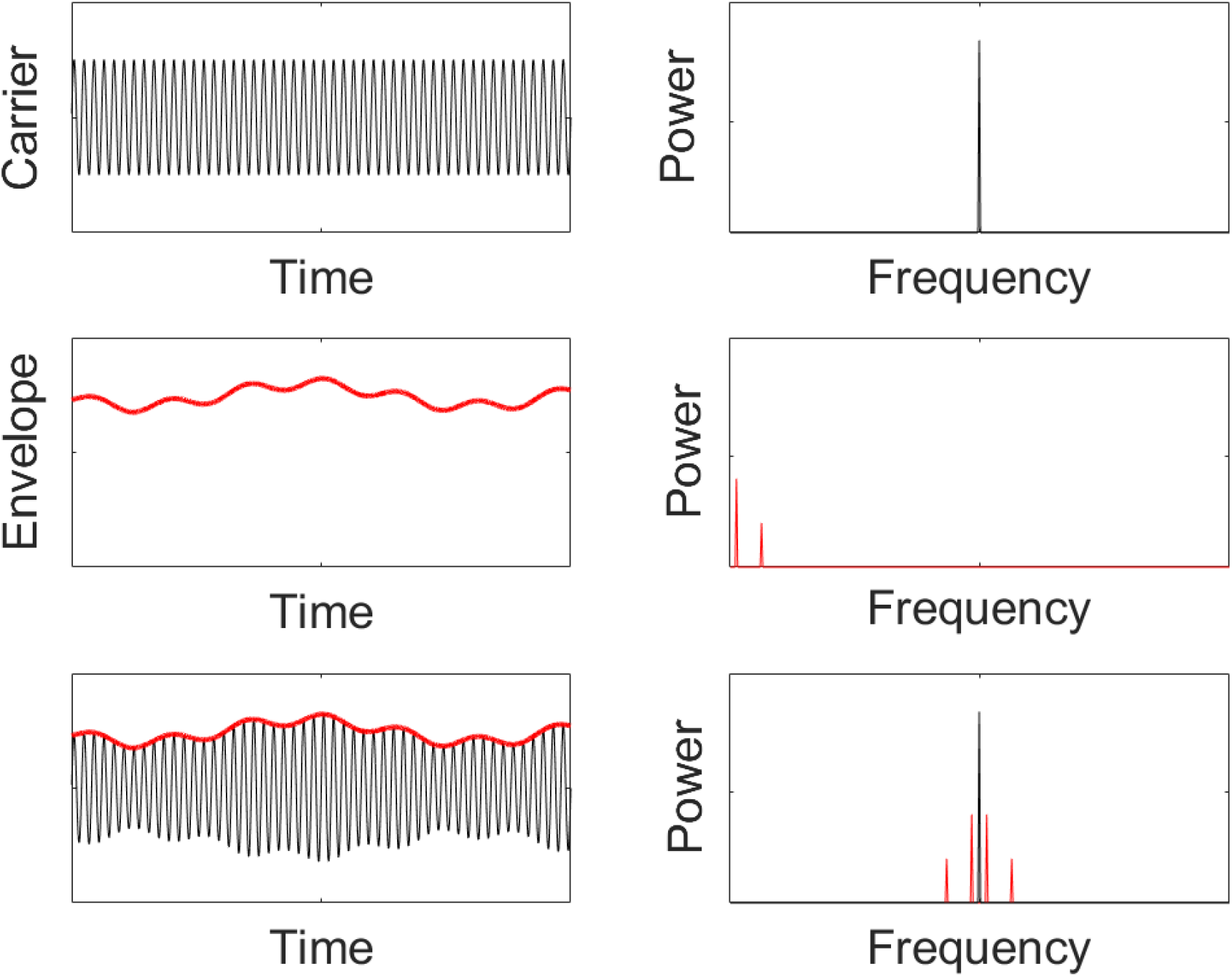
Spectral symmetry is a consequence of amplitude modulation. A pure sinusoidal carrier signal (top) with some amplitude modulation (middle, envelope) results in a signal with a symmetrical power spectrum (bottom). The side-peaks occur around the carrier frequency (black), plus and minus the frequencies of the envelope modulation (red). If the idealised voltage during tACS has its amplitude modulated by rhythmic changes in impedance (remembering that the current is controlled), this should result in spectral peaks around the frequency of tACS equal to the rhythmicity observed in the voltage envelope.

While it has been suggested that mechanical failure and regularisation during the beamforming pipeline may be the cause of this spectral symmetry (Neuling et al., 2017; see also Bland, 2019), the physics underlying tACS artefact modulations is well understood and suggests instead a predominantly physiological origin (Noury and Siegel, 2018). Since transcranial *current* stimulation techniques are current-controlled, Ohm’s law dictates that any change in body impedance will change the stimulation voltage. Since heartbeat and respiration both rhythmically modulate body impedance (Noury et al., 2016), the voltage will vary with these rhythms. The impedances under the EEG electrodes are also time-varying, as they too make contact with the scalp. In the MEG, the movement of head position due to ballistocardiographic and respiratory effort leads to rhythmic modulations in the magnetic field, further contributing to the time-varying nature of tACS artefacts. Amplitude modulations by heartbeat have also been detected in the MEG during transcranial direct current stimulation (tDCS, Marshall et al., 2016), contaminating slow oscillations (i.e., ‘symmetrically’ around 0 Hz, Noury et al., 2016).

Since the time-varying nature of tACS (and indeed tDCS) has a physiological origin, resultant artefacts are not perfectly correlated across sensors (as would be the case if each sensor contained a scaled version of the current). Therefore, any method that relies on the decomposition of the M/EEG covariance matrix (e.g., PCA) will fail to fully capture these artefacts. Indeed, the beamformer (which effectively suppresses correlated noise sources, Neuling et al., 2015; Mäkelä et al., 2017) fails because of imperfect collinearity between tACS artefacts at the sensor level (Noury and Siegel, 2018). Therefore, any approach designed to salvage an uncontaminated spectrum of endogenous activity during tACS first requires methods of artefact modelling to take into account the time-varying nature of tACS artefacts across M/EEG sensors. This is necessary to make a valid comparison with a control spectrum observed during sham (where no stimulation is applied and the side-peaks are absent), or during tACS applied at a different frequency (where the spectral symmetry occurs elsewhere in the frequency domain).

A good starting place for the development of time-varying models of tACS artefacts is the analytic signal (i.e., with its Hilbert transform), exploiting each sensor’s instantaneous amplitude and phase. As shown in **Figure 3**, valuable information about amplitude modulations by heartbeat and respiration is resolvable from the envelope of the tACS artefact. However, variability in heart rate and breath rate, and the non-sinusoidal nature of these perturbations, introduces harmonics in the envelope which can contaminate the M/EEG spectrum by as much as ±10 Hz from the frequency of tACS. Following the characterisation of the transfer function between the tACS current and its artefact (Noury and Siegel, 2017), here I show how a time-varying approach to modelling artefacts of tACS in the M/EEG—with consideration of the nonlinear modulations by heartbeat and respiration—can dramatically suppress artefactual side-peaks in the online M/EEG spectrum. I also discuss the feasibility of using this same approach in the absence of concurrent measurements of heartbeat and respiration, as occurs in most studies that combine non-invasive brain stimulation with neuroimaging.

**Figure 3.**
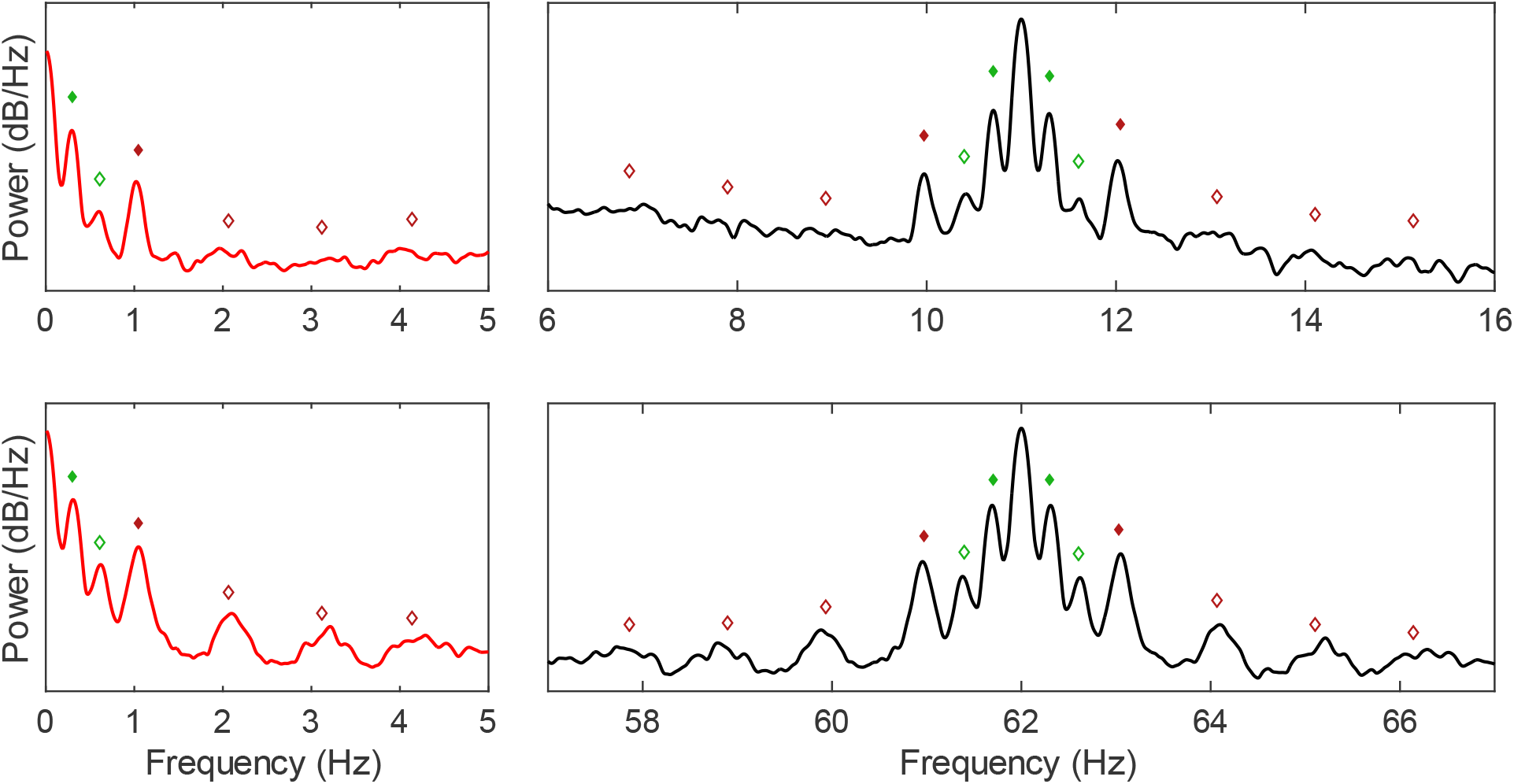
The artefact envelope captures amplitude modulations by heartbeat and respiration. The power spectrum of the tACS envelope (red, left) captures nonlinear amplitude modulations by both heartbeat (red diamonds) and respiration (green diamonds), contaminating the spectrum (black, right) symmetrically around the frequency of tACS (11 Hz, top; 62 Hz, bottom). The harmonics of heart rate and breath rate are also detectable (unfilled diamonds). These power spectra (EEG sensor PO10 displayed) were averaged across 12 available periods of 11 Hz and 62 Hz tACS for one representative participant.

## METHOD AND RESULTS

### Data

The data used here were reported in three previous publications (Noury et al., 2016; Noury and Siegel, 2017, 2018), the first of which contains detailed descriptions of the hardware, experiment setup, and preprocessing steps. Data were collected from 4 healthy male participants across 6 experimental runs, with 72-channel EEG and 272-channel MEG simultaneously recorded throughout. Each experimental run comprised two applications of 11 Hz, 62 Hz, and sham tACS— each lasting 66 seconds (a total of 12 applications per tACS condition). Stimulation current was applied at 1 mA peak-to-peak between right occipital (O10) and right parietal (CP4) electrodes (**Figure 1B**, black). The EEG system was used to record the injected current (voltage drop across a 200 Ohm resistor in series to the head) and the electrocardiogram. Respiration was recorded with a piezoelectric belt transducer.

### Preprocessing

M/EEG data were highpass filtered down to 0.25 Hz, lowpass filtered up to 90 Hz, and notch filtered from 48.8 Hz to 50.2 Hz (to remove 50 Hz mains power). All were 6^th^-order zero-phase Butterworth filters. EEG data were downsampled to 1000 Hz from 10000 Hz, and MEG data were downsampled to 781.25 Hz from 2343.75 Hz. To account for filtering edge effects, the first and last 1.5s were trimmed from the recordings. A version of the artefactual M/EEG was further bandpass filtered between tACS ± 5 Hz with a 6^th^-order zero-phase Butterworth filter for computing the instantaneous amplitude and phase.

### A Fixed Model

Artefacts of tACS can be crudely approximated using a fixed (time-invariant) sinusoidal template (Bland and Sale, 2019). This section describes a computationally efficient method to linearly fit a fixed template to artefactual M/EEG data, performed separately for each sensor. While this alone does a poor job of capturing tACS artefacts (because the artefacts are time-varying), rhythmic changes in amplitude and phase can be added to this basis template, thus transforming it into a time-varying model as required.

Given data *d* of length *n*, form column vectors for the artefactual recording *s* over time *t* given the sampling frequency *Fs*.

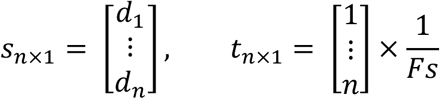

Form a predictor matrix over time with columns for the bias (all ones), and with unit sine and cosine components (used together for amplitude and phase), given some fundamental frequency *F* (e.g., the claimed tACS frequency).

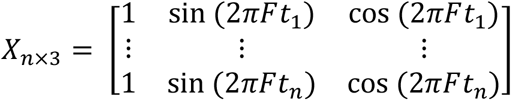

Use ordinary least squares to solve the normal equations for the parameters *θ*.^1^ A desirable property of this approach is that the Gramian matrix *X*^*T*^*X* remains 3 × 3 irrespective of the length of *s*, keeping computation time for the inverse low. Iterating this procedure over *F* converges quickly upon the optimal frequency, especially since the claimed tACS frequency makes a great seed (e.g., for the Nelder–Mead simplex algorithm; fminsearch). The fixed sinusoidal model is ŝ (i.e., sharing the same dimensions as *s*_*n*×1_). *T* denotes the matrix transpose.

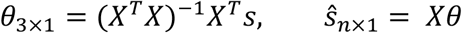

Equivalently, the model ŝ has bias *b*, amplitude *α*, and phase offset *ϕ*. These constants can be reconstructed from the parameters *θ*, where atan2(*y, x*) computes the four-quadrant inverse tangent—the angle between the positive *x*-axis and the vector representing (*x, y*) in the Cartesian plane.

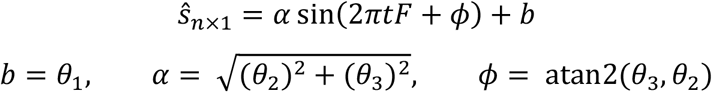

With the fixed model estimated, this time-invariant sinusoid (*b, α, ϕ* all constants) would form the carrier for any amplitude and phase modulations introduced at the known times of any heart- and respiration-related perturbations, thus transforming it into a time-varying template.

### Estimating Frequency

The frequency of tACS is ideally estimated from recordings of its current output, free from endogenous brain activity (Noury et al., 2016). In order to fit good sinusoidal templates to the artefacts, the frequency needs to be known with great precision (e.g., in the microhertz range). Here, this was achieved by seeding a Nelder–Mead simplex at the claimed tACS frequency and converging on a least squares minimum across the full recording of the current output, while allowing amplitude, phase, and any baseline (bias) to vary freely (stopping tolerance set at 10^−12^ Hz for frequency). In the EEG, this resulted in standard deviations of 0.09 and 0.52 μHz for 11 Hz and 62 Hz tACS, respectively. Similar precision was achieved previously by estimating frequency independently across halved segments of the current output (Noury et al., 2016).

If no recording of the current output is available, high precision can still be achieved using the artefactual M/EEG (especially in sensors where the tACS amplitude is largest). Generally speaking, sensors with artefact amplitudes within the top 50%–10% shared similar variation in frequency estimates with the current output recordings discussed above. Similar performance can also be achieved using the per-sensor frequency, though any differences between sensors are necessarily errors (since the frequency of tACS is constant across sensors). I speculate that the inclusion of harmonics in the simplex may improve accuracy of frequency estimates, though I have not done so here because the current output recordings were available.

### A Time-Varying Model

The basis template derived from the fixed model approach will fail to capture any time-varying changes in the amplitude and phase of tACS artefacts. However, these modulations can be estimated from the artefactual M/EEG. Consider the recording from a single M/EEG sensor *s* as a summation of artefact *A* and artefact-free activity—or brain activity, *b*—over time.

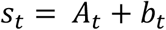

If the artefact *A* were known, it would be trivial to subtract it from s. However, *A* is typically unknown, and so estimates of its instantaneous amplitude and phase must instead be derived from *s*. This is potentially problematic because endogenous activity *b* will therefore contribute to these estimates (Noury and Siegel, 2017). However, since the tACS artefact dominates the recording, this nevertheless provides a good starting place for estimating *A*. The recording *s* will contain modulations by heartbeat and respiration in its instantaneous amplitude and phase, which can be derived from its analytic signal, and these modulations should be uncorrelated with ongoing endogenous activity.

The analytic signal has real part *s* and imaginary part *ℌ*{*s*}, its Hilbert transform. The instantaneous amplitude *α* is the complex magnitude of the analytic signal; the instantaneous phase *ϕ* is the argument. The original recording *s* can be reconstructed from these, where *Re* takes only the real part. Note that *α* and *ϕ* were previously constants, but are now time-varying.

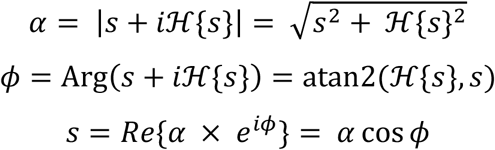

To improve estimates of the instantaneous amplitude and phase, the artefact can be better isolated from ongoing brain activity by bandpass filtering the contaminated M/EEG around the frequency of stimulation. Alternatively, the envelope of the recording can be lowpass filtered, suppressing low-frequency amplitude modulations (**Figure 4**). However, this fails to *selectively* suppress only the physiological amplitude modulations, and will tend to overfit, resulting in excessively dampened power around the frequency of tACS. Ideally, we wish only to suppress the nuisance modulations, and to do this both for amplitude and phase.

**Figure 4.**
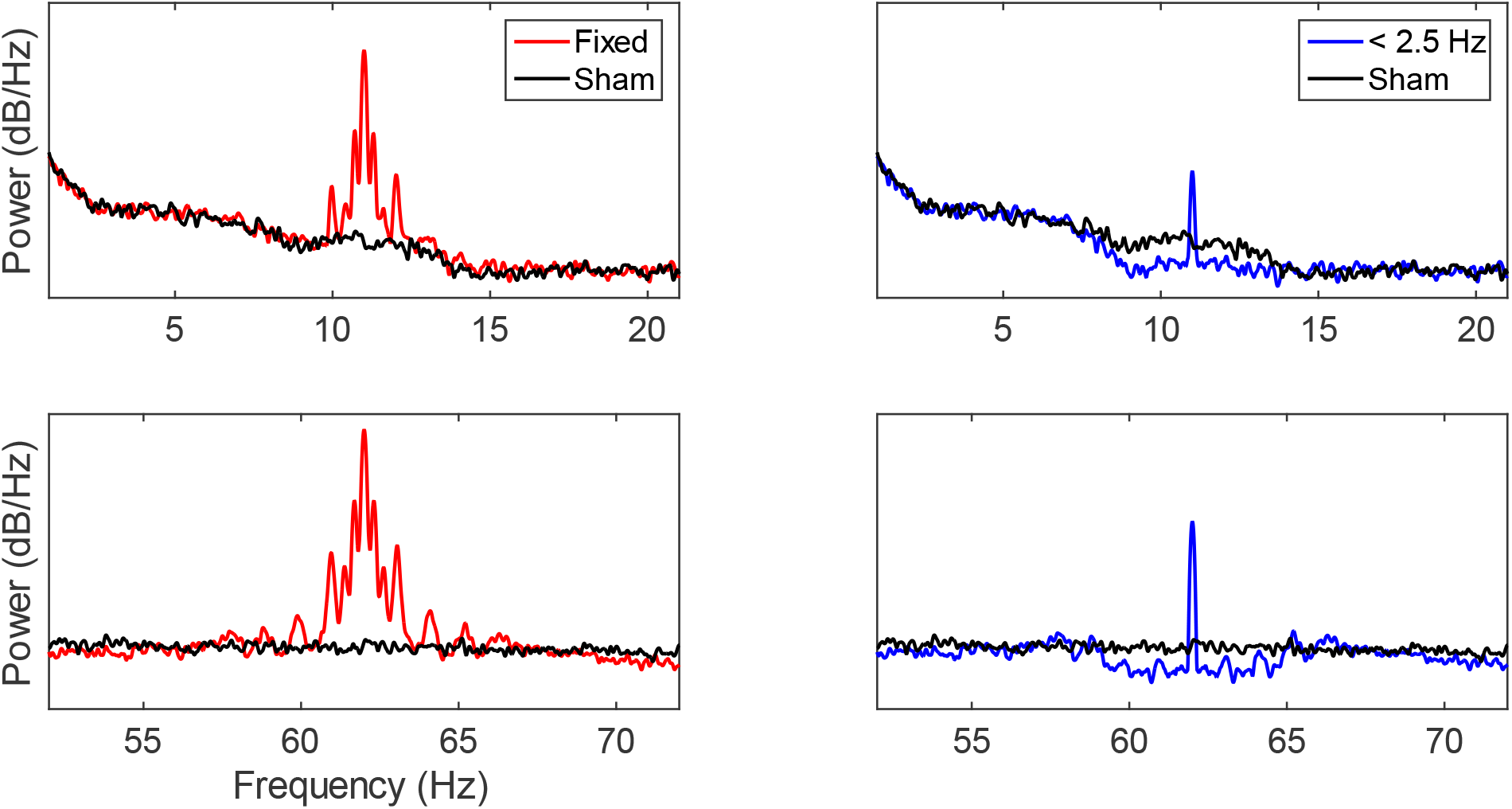
The lowpass envelope non-selectively suppresses side-peaks. Compared to a fixed-amplitude template subtraction approach (red), lowpass filtering the envelope (with stopband at 2.5 Hz, blue) and using this as the amplitude of the time-varying template dampens the spectral symmetry around the frequency of tACS (11 Hz, top; 62 Hz, bottom). However, since all frequency content below 2.5 Hz is preserved, this method does not suppress the nuisance side-peaks selectively, and tends to over-dampen the spectra around the carrier frequency (that of tACS) compared with the spectra observed during sham tACS (black). These power spectra (EEG sensor PO10 displayed) were averaged across 12 available periods of 11 Hz and 62 Hz tACS for one representative participant.

### Time-Locked Perturbations

Rather than indiscriminately including all low-frequency amplitude modulations observed below some stopband (e.g., 2.5 Hz; **Figure 4**), a better approach would be to *selectively* suppress both amplitude and phase modulations by heartbeat and respiration. Since the electrocardiogram and the belt transducer provide traces for these physiological rhythms, time-locked models of their effects can be formed from the timing of events observed in these recordings. Here, the timing of heartbeats and points of inspiration can be estimated from the peaks observed in the electrocardiogram and by the belt transducer. Following the same principle for forming event-related potentials, these known event times are used to index into the instantaneous amplitude (envelope) to form the mean-removed time-locked amplitude modulations by heartbeat and respiration (e.g., windowed over three cycles; **Figure 5**). A similar procedure can be performed for the phase modulation, though the instantaneous phase must first be unwrapped (linearised) and detrended. If brain activity is uncorrelated with heartbeat and respiration, the time-locked models will tend to only capture heartbeat- and respiration-related perturbations to amplitude and phase. The time-locked perturbations to the amplitude and phase by heartbeat and respiration can now be added to the otherwise fixed sinusoidal template based on the same event timing used to form the event-locked models. The resultant time-varying template selectively suppresses these physiologically derived side-peaks and their harmonics (**Figure 6**).

**Figure 5.**
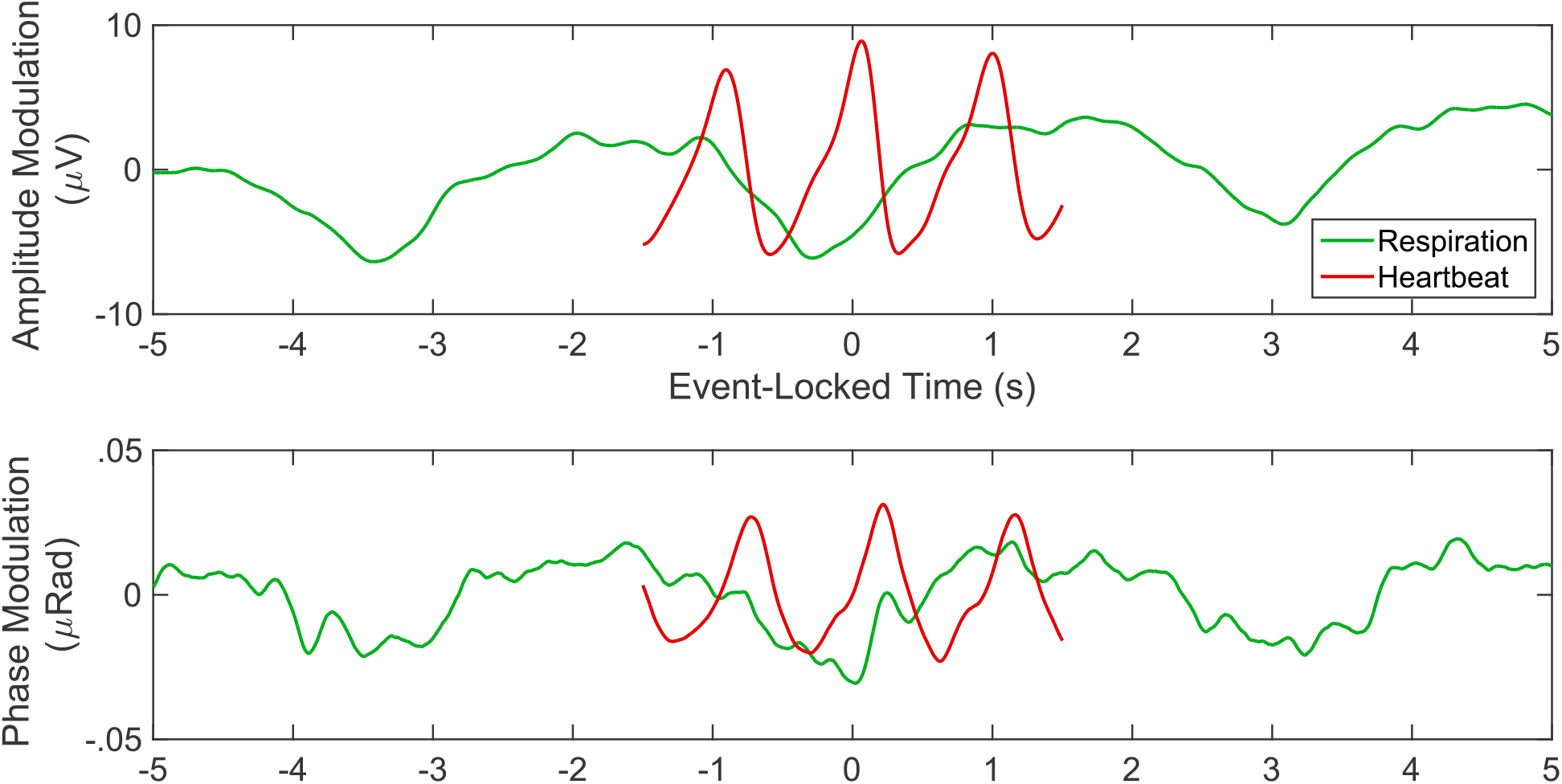
Event-related perturbations to amplitude and phase by heartbeat and respiration. Time-locked models of the mean-removed heartbeat- and respiration-related modulations to amplitude and phase can be formed based on the known event times in the simultaneously recorded electrocardiogram and respiration data. The amplitude modulation is taken from the envelope (instantaneous amplitude); the phase modulation is taken from the unwrapped and detrended instantaneous phase. These event-related perturbations (EEG sensor O9 displayed) were averaged across 12 available periods of 62 Hz tACS for one representative participant. Note that the modulations themselves are subtle—the average amplitude of the tACS artefact at O9 was 16.37 mV for this participant (amplitude modulations represent approximately .0005% of the tACS artefact amplitude). However, these effects remain within the range of biologically plausible endogenous activity because tACS artefacts are so large.

**Figure 6.**
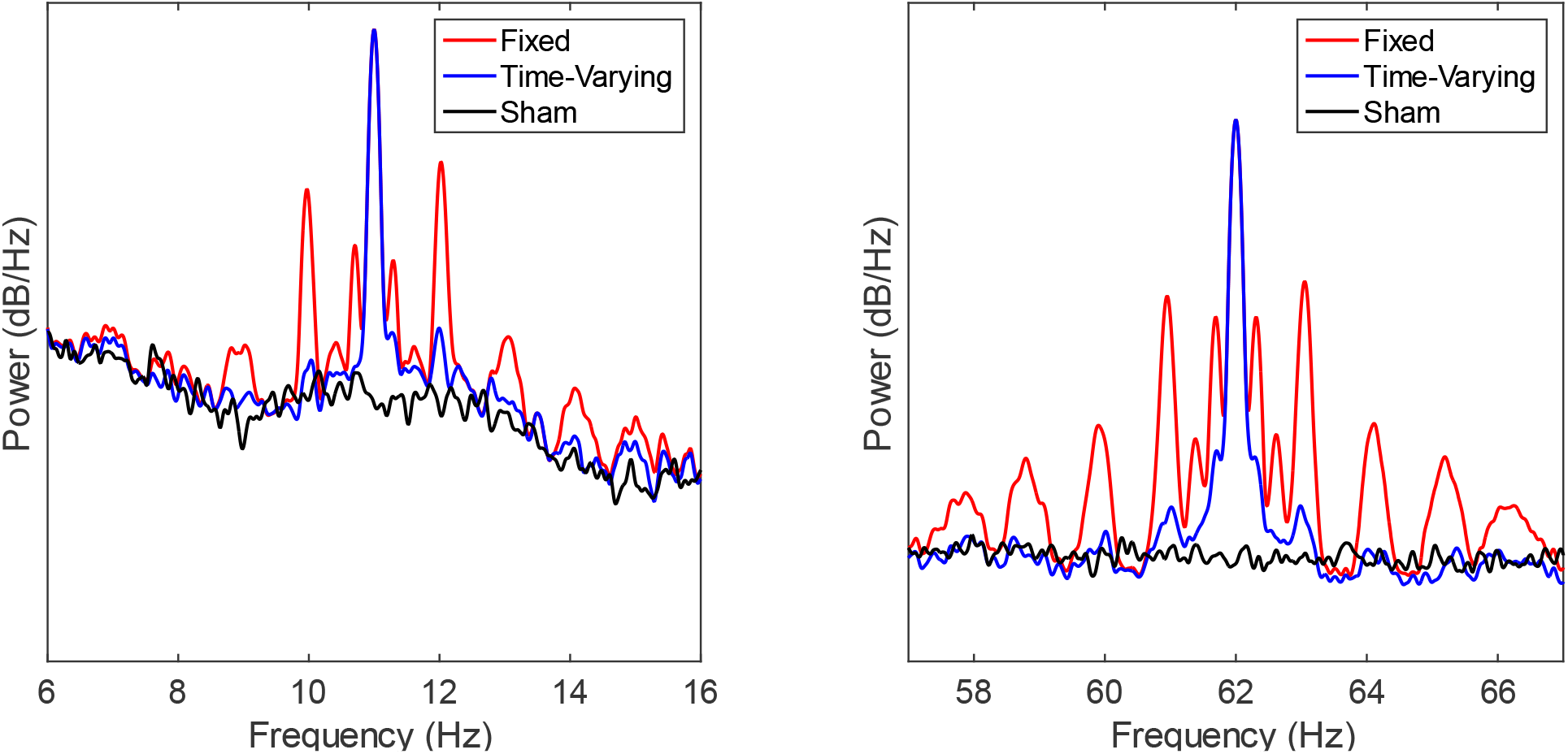
A time-varying model selectively suppresses the spectral symmetry. Compared to a fixed-amplitude template subtraction approach (red), the model with variable amplitude and phase determined by the captured time-locked perturbations (blue) dampens the spectral symmetry around the frequency of tACS (11 Hz, left; 62 Hz, right). Note that a large peak around the fundamental frequency (that of tACS) remains even with a time-varying model, likely reflecting ultraslow amplitude modulations (e.g., that of electrode impedance over time)—though potentially also contains true entrainment by tACS. These power spectra (EEG sensor O9 displayed) were averaged across 12 available periods of 11 Hz and 62 Hz tACS for one representative participant.

### Naïve Heartbeat- and Respiration-Locked Timing

Since heartbeat and respiration tend to be rhythmic—and this rhythmicity is identifiable in the spectrum of the artefactual M/EEG envelope—event-locked heartbeat and respiration models may be estimated without concurrent recordings of heartbeat and respiration (i.e., in cases where the true event timing is unknown). While the effects of heartbeat and respiration on amplitude and phase appear largely consistent over successive events, recent work suggests variable timing at least between events in the electrocardiogram and resultant ballistocardiographic artefacts (Marino et al., 2018a). A better approach may therefore be to use points of maximal cross-correlation (Marino et al., 2018b) to self-generate (estimate) the timing of heartbeat- and respiration-related amplitude modulations, rather than from the events observed in the physiological traces. The timings yielded from such an approach could then be used to index into the instantaneous phase, since the phase effects are themselves much subtler (Noury and Siegel, 2017).

Based on the approximate heart rate and breath rate (which can be yielded reliably from the envelope spectra; **Figure 3**), adjacent windows can be seeded across the envelope to self-generate initial heartbeat- and respiration-related amplitude modulation (i.e., the window average). A nice property of this approach is that it makes no assumption about the form of the amplitude modulations at each sensor. Variability in heart rate and breath rate over time makes these initial models generally dampened approximations (**Figure 7**), but they can still be used to find the points of maximal cross-correlation between the window average and each individual window. These times can then be used to iteratively update the window average and quickly shift the windows toward the points of maximal cross-correlation. After just a few iterations, this method self-generates the original model formed by the true event-timings (**Figure 7**; i.e., generated from the times sourced from the actual electrocardiogram). Alternatively, since the time-locked models appear quite similar in shape across sensors and participants, cross-correlations may be performed between a previously generated model and each window (though this will of course self-select for event-timings that approximate this assumed model, which may not be valid for some sensors or participants).

**Figure 7.**
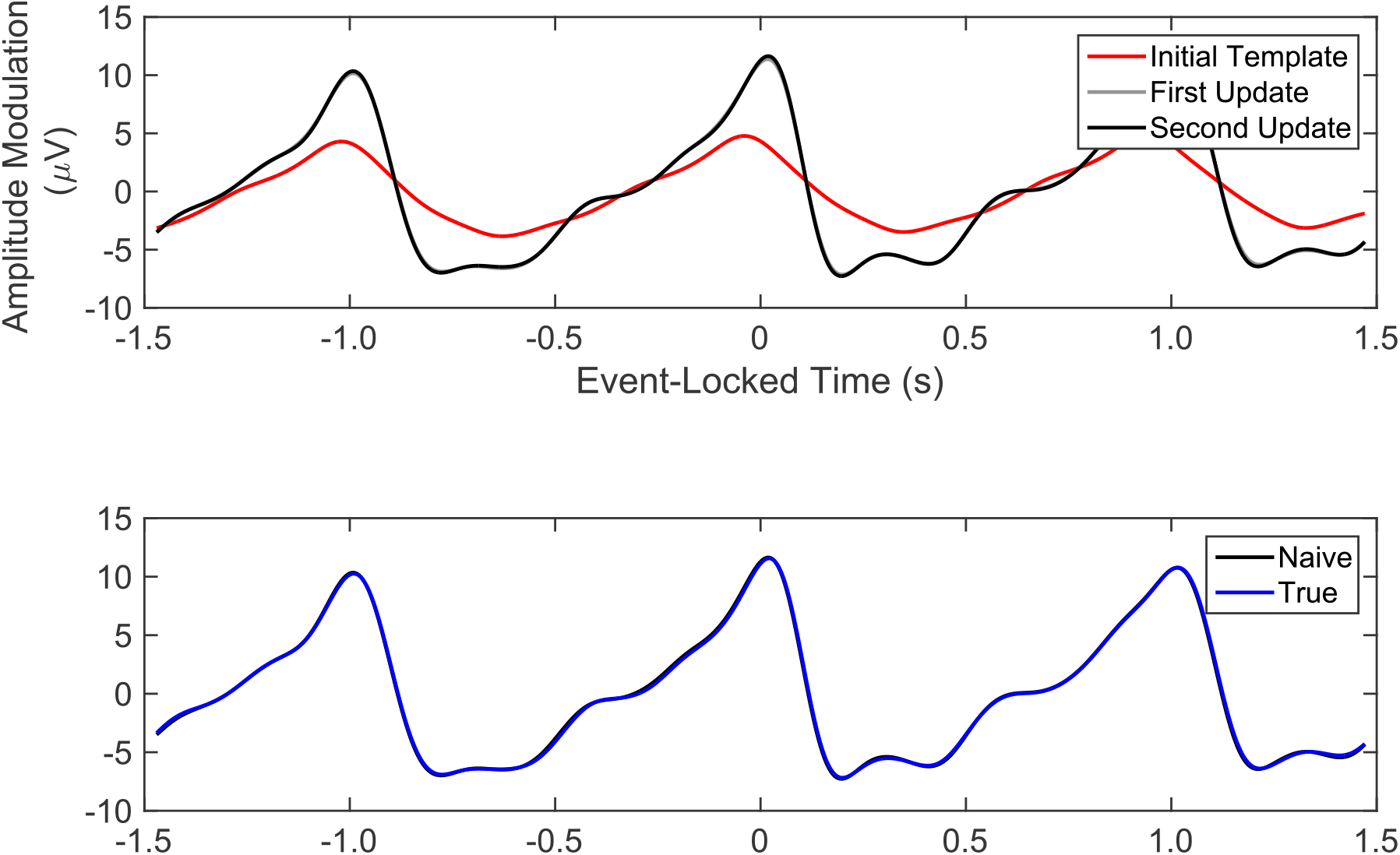
Naively generated model of the event-related perturbation to amplitude by heartbeat. The initial window average (red)—seeded based on the estimated heart rate from the envelope—forms a dampened approximation of the true heartbeat modulation. By iteratively updating the windows based on the point of maximum cross-correlation with the window average, a naïve model is quickly self-generated in the absence of the known event timing (i.e., where the model is naïve to the electrocardiogram). This naive model (shown after just two iterations, black; first iteration largely hidden underneath, grey) very closely approximates the true heartbeat-related amplitude modulation (blue), which is formed based on the known heartbeat times in the electrocardiogram. These self-generated event-times can also be used to index into the instantaneous phase to form phase modulations (not shown).

## DISCUSSION

Online analysis of concurrent tACS–M/EEG requires robust and effective approaches for suppressing or removing artefacts of stimulation. Without fully accounting for the spectral symmetry introduced by heartbeat- and respiration-related modulations to the tACS amplitude and phase, side-peaks may contribute spuriously to any evidence for neural entrainment, particularly when comparing to a spectrum observed during sham stimulation (where these side-peaks are absent) or to a spectrum observed with another frequency of tACS (where spectral symmetry occurs elsewhere in the frequency domain). The wide range of published approaches for removing artefacts of tACS from M/EEG illustrates how difficult a problem this is. However, existing approaches have universally assumed tACS artefacts are either time-invariant or space-invariant (i.e., perfectly collinear over sensors).

Here, I demonstrate the feasibility of linearly solving base templates for tACS artefacts and subsequently forming time-varying models by introducing event-locked heartbeat-and respiration-related perturbations to the amplitude and phase of tACS. I also demonstrate that the known timing of heartbeats and inspiration can be used to index into the envelope (amplitude modulations) and instantaneous phase (phase modulations) of the M/EEG on a per-sensor basis, assuming neither time- nor sensor-invariance. Introducing these event-locked perturbations to an otherwise fixed sinusoidal model dramatically suppresses the spectral symmetry observed in the recovered spectrum, and provides a proof of principle for removing nonlinear modulations to tACS from the M/EEG. Further, the amplitude modulations by heartbeat and respiration are so large that the timing of these events could be estimated from the artefactual M/EEG. Therefore, I conclude that time-varying approaches are feasible even in the absence of simultaneously recorded heartbeat and respiration. This would provide researchers a means to analyse their online M/EEG even when these physiological measures have not been recorded.

### Limitations and Future Directions

While heartbeat- and respiration-related artefacts can be modelled and suppressed in the M/EEG, this assumes the modulations themselves remain stable over time. However, infra-slow (< 0.2 Hz) amplitude modulations can be introduced by impedance changes under the tACS pads (and under the EEG electrodes), which means the extent to which heartbeat and respiration modulate tACS artefacts is itself time-varying (i.e., the modulations will be largest when the tACS artefact is largest)—the periodic and aperiodic modulations interact (see Bland and Sale, 2019). For example, if impedance under the tACS pads decreases over time, the tACS voltage and corresponding artefact will become smaller and so too will the modulations to it by heartbeat and respiration.

These infra-slow amplitude modulations are a likely cause of the discernible peaks that remain around the tACS frequency, even after suppressing the side-peaks. This could be addressed in future by modelling amplitude modulations as a percentage of the tACS amplitude, so that slower changes to the amplitude (say by impedance) also scale its amplitude modulations by heartbeat and respiration. These infra-slow amplitude modulations can be estimated by smoothing the instantaneous amplitude, lowpass filtering it up to 0.2 Hz, or by fitting an ideal sinusoid within a sliding window across the recording and evaluating the point-wise amplitude. Low-frequency tACS (e.g., 0.1–5 Hz) may further complicate modelling slow amplitude- and phase-modulations.

For most complex tACS waveforms (i.e., not mono-sinusoidal), time-varying approaches based on event-locked perturbations are not feasible since the instantaneous amplitude and phase are not constant in the absence of nonlinear modulations. For example, piecewise linear (sawtooth) tACS (Dowsett and Herrmann, 2016) will still be susceptible to the same nonlinear modulations, but these modulations will depend heavily on the phase-cycle of the waveform (e.g., amplitude modulations would be largest toward the peak of the sawtooth). Instead, beamforming—which will suppress a collinear artefact irrespective of its form—may be a better approach, though will of course also fail to remove nonlinear modulations to amplitude and phase. Indeed, even mono-sinuoidal tACS has harmonics at multiples of the fundamental. While these harmonic components are comparably small, spectral symmetry may plausibly occur around multiples of the tACS frequency, though to a much lesser extent than around the fundamental.

Self-generating a model of the amplitude modulation by respiration can be complicated when it has a harmonic near the heart rate (e.g., 0.33 Hz breath rate and 1 Hz heart rate). In these cases, the breath-related model may spuriously capture the nested heartbeat-related perturbations. If this occurs, removing both event-locked models from the envelope may plausibly “double dip” on the heartbeat-related modulation, effectively re-introducing it to the envelope. This may be why some small residual heart-related symmetry remains in the recovered spectra (**Figure 6**). To minimise this problem, the heartbeat-related amplitude modulations may first be removed from the tACS envelope before computing the breath-related amplitude modulation based on the belt transducer (and especially if computing naive breath-related amplitude modulation using the cross-correlations). I speculate that singular spectrum analysis of the envelope may provide an alternative method for approximating the event-timings of heartbeat-and respiration-related modulations while remaining naïve to their form.

The field of tACS research has benefitted greatly from amplitude-modulated tACS (Witkowski et al., 2016; Negahbani et al., 2018; Haslacher et al., 2021), where the carrier frequency (usually a very high frequency; e.g., 220 Hz) is purposely amplitude modulated at the frequency of physiological interest (e.g., 10 Hz). If the modulation frequency (envelope) can itself reliably entrain neural activity at that frequency, then the problem of nonlinear modulations will be shifted into the higher frequency domain, and will instead occur symmetrically around the carrier, and symmetrically around the carrier plus-and-minus the modulation frequency. In principle, this would effectively avoid any spectral overlap between the brain signals of interest and the artefact of tACS. However, some work suggests the M/EEG spectrum can remain corrupted near the modulation frequency (Minami and Amano, 2017; Kasten et al., 2018), though to a much lesser extent than observed during mono-sinusoidal tACS.

## Conclusion

Time-varying approaches for modelling artefacts of tACS are necessary for capturing the nonlinear modulations that contaminate concurrent M/EEG. Improvements in our ability to capture artefacts of tACS (e.g., recording the voltage output of tACS) and the timing of physiologically-derived modulations (i.e., that of heartbeat and respiration) will pave the way to sound online spectral analysis. The future validation of these time-varying approaches on both real and fabricated M/EEG data is a critical step before claiming online evidence for entrainment by tACS.

## ACKNOWLEDGEMENTS

I thank Nima Noury, Joerg Hipp, and Markus Siegel for making their original data available. The original data were collected under support by the Centre for Integrative Neuroscience (Deutsche Forschungsgemeinschaft, EXC 307). I thank Martin Sale and Brendan Keane for providing comments on an earlier version of this manuscript.

## Footnotes

1 This process can be adapted to include any harmonics of *F* (or indeed any frequencies one wishes by concatenating the columns of *X* with unit sine and cosine components for every frequency of interest (i.e., the Vaníček method of least squares spectral analysis; Vaníček, 1969).

## Notes

**CONFLICT OF INTEREST STATEMENT** The author declares that the research was conducted in the absence of any commercial or financial relationships that could be construed as a potential conflict of interest.

### Competing Interest Statement

The authors have declared no competing interest.

## REFERENCES

Bergmann, T. O., Karabanov, A., Hartwigsen, G., Thielscher, A., & Siebner, H. R. (2016). Combining non-invasive transcranial brain stimulation with neuroimaging and electrophysiology: Current approaches and future perspectives. NeuroImage, 140, 4–19.

Bland, N. S. (2019). Oscillating neural networks: Perspectives from rhythmic brain stimulation. The University of Queensland, Australia.

Bland, N. S., Mattingley, J. B., & Sale, M. V. (2018). No evidence for phase-specific effects of 40 Hz HD–tACS on multiple object tracking. Frontiers in Psychology, 9, 304.

Bland, N. S., Mattingley, J. B., & Sale, M. V. (2020). Gamma coherence mediates interhemispheric integration during multiple object tracking. Journal of Neurophysiology, 123(5), 1630–1644.

Bland, N. S., & Sale, M. V. (2019). Current challenges: The ups and downs of tACS. Experimental Brain Research, 237(12), 3071–3088.

Fiene, M., Schwab, B. C., Misselhorn, J., Herrmann, C. S., Schneider, T. R., & Engel, A. K. (2020). Phase-specific manipulation of rhythmic brain activity by transcranial alternating current stimulation. Brain Stimulation, 13(5), 1254–1262.

Geffen, A., Bland, N., & Sale, M. V. (2021). Effects of slow oscillatory transcranial alternating current stimulation on motor cortical excitability assessed by transcranial magnetic stimulation. bioRxiv. doi: 10.1101/2021.05.13.444101

Haslacher, D., Nasr, K., Robinson, S. E., Braun, C., & Soekadar, S. R. (2021). Stimulation Artifact Source Separation (SASS) for assessing electric brain oscillations during transcranial alternating current stimulation (tACS). NeuroImage, 228, 117571.

Helfrich, R. F., Schneider, T. R., Rach, S., Trautmann-Lengsfeld, S. A., Engel, A. K., & Herrmann, C. S. (2014). Entrainment of brain oscillations by transcranial alternating current stimulation. Current Biology, 24(3), 333–339.

Herrmann, C. S., Rach, S., Neuling, T., & Strüber, D. (2013). Transcranial alternating current stimulation: A review of the underlying mechanisms and modulation of cognitive processes. Frontiers in Human Neuroscience, 7:279.

Herrmann, C. S., & Strüber, D. (2017). What can transcranial alternating current stimulation tell us about brain oscillations?. Current Behavioral Neuroscience Reports, 4(2), 128–137.

Huang, W. A., Stitt, I. M., Negahbani, E., Passey, D. J., Ahn, S., Davey, M., … & Fröhlich, F. (2021). Transcranial alternating current stimulation entrains alpha oscillations by preferential phase synchronization of fast-spiking cortical neurons to stimulation waveform. Nature Communications, 12(1), 1–20.

Kasten, F. H., & Herrmann, C. S. (2019). Recovering brain dynamics during concurrent tACS-M/EEG: an overview of analysis approaches and their methodological and interpretational pitfalls. Brain Topography, 32(6), 1013–1019.

Kasten, F. H., Negahbani, E., Fröhlich, F., & Herrmann, C. S. (2018). Non-linear transfer characteristics of stimulation and recording hardware account for spurious low-frequency artifacts during amplitude modulated transcranial alternating current stimulation (AM-tACS). NeuroImage, 179, 134–143.

Kohli, S., & Casson, A. J. (2015). Removal of transcranial ac current stimulation artifact from simultaneous EEG recordings by superposition of moving averages. In Engineering in Medicine and Biology Society (EMBC), 37th Annual International Conference of the IEEE (pp. 3436–3439).

Kohli, S., & Casson, A. J. (2019). Removal of gross artifacts of transcranial alternating current stimulation in simultaneous EEG monitoring. Sensors, 19(1), 190.

Mäkelä, N., Sarvas, J., & Ilmoniemi, R. J. (2017). A simple reason why beamformer may (not) remove the tACS-induced artifact in MEG. Brain Stimulation: Basic, Translational, and Clinical Research in Neuromodulation, 10(4), e66–e67.

Marino, M., Liu, Q., Del Castello, M., Corsi, C., Wenderoth, N., & Mantini, D. (2018a). Heart– Brain interactions in the MR environment: Characterization of the ballistocardiogram in EEG signals collected during simultaneous fMRI. Brain Topography, 31(3), 337–345.

Marino, M., Liu, Q., Koudelka, V., Porcaro, C., Hlinka, J., Wenderoth, N., & Mantini, D. (2018b). Adaptive optimal basis set for BCG artifact removal in simultaneous EEG-fMRI. Scientific Reports, 8(1), 8902.

Marshall, T. R., Esterer, S., Herring, J. D., Bergmann, T. O., & Jensen, O. (2016). On the relationship between cortical excitability and visual oscillatory responses—A concurrent tDCS–MEG study. Neuroimage, 140, 41–49.

Minami, S., & Amano, K. (2017). Illusory jitter perceived at the frequency of alpha oscillations. Current Biology, 27(15), 2344–2351.

Negahbani, E., Kasten, F. H., Herrmann, C. S., & Fröhlich, F. (2018). Targeting alpha-band oscillations in a cortical model with amplitude-modulated high-frequency transcranial electric stimulation. Neuroimage, 173, 3–12.

Neuling, T., Ruhnau, P., Fuscà, M., Demarchi, G., Herrmann, C. S., & Weisz, N. (2015). Friends, not foes: Magnetoencephalography as a tool to uncover brain dynamics during transcranial alternating current stimulation. NeuroImage, 118, 406–413.

Neuling, T., Ruhnau, P., Weisz, N., Herrmann, C. S., & Demarchi, G. (2017). Faith and oscillations recovered: On analyzing EEG/MEG signals during tACS. Neuroimage, 147, 960–963.

Noury, N., Hipp, J. F., & Siegel, M. (2016). Physiological processes non-linearly affect electrophysiological recordings during transcranial electric stimulation. NeuroImage, 140, 99–109.

Noury, N., & Siegel, M. (2017). Phase properties of transcranial electrical stimulation artifacts in electrophysiological recordings. NeuroImage, 158, 406–416.

Noury, N., & Siegel, M. (2018). Analyzing EEG and MEG signals recorded during tES, a reply. Neuroimage, 167, 53–61.

Polanía, R., Nitsche, M. A., & Ruff, C. C. (2018). Studying and modifying brain function with non-invasive brain stimulation. Nature Neuroscience, 21, 174–187.

Santos Monteiro, T, Heise, K., Swinnen, S., & Mantini, D. (2015). A novel approach for the removal of tACS artifacts from high-density EEG recordings. Frontiers in Neuroinformatics. Conference Abstract: Second Belgian Neuroinformatics Congress.

Soekadar, S.R., Herring, J.D., & McGonigle, D. (2016). Transcranial electric stimulation (tES) and NeuroImaging: The state-of-the-art, new insights and prospects in basic and clinical neuroscience. NeuroImage, 140, 1–3.

Vaníček, P. (1969). Approximate spectral analysis by least-squares fit. Astrophysics and Space Science, 4(4), 387–391.

Veniero, D., Vossen, A., Gross, J., & Thut, G. (2015). Lasting EEG/MEG aftereffects of rhythmic transcranial brain stimulation: level of control over oscillatory network activity. Frontiers in Cellular Neuroscience, 9:477.

Vieira, P. G., Krause, M. R., & Pack, C. C. (2020). tACS entrains neural activity while somatosensory input is blocked. PLoS Biology, 18(10), e3000834.

Voss, U., Holzmann, R., Hobson, A., Paulus, W., Koppehele-Gossel, J., Klimke, A., & Nitsche, M. A. (2014). Induction of self awareness in dreams through frontal low current stimulation of gamma activity. Nature Neuroscience, 17(6), 810–812.

Witkowski, M., Garcia-Cossio, E., Chander, B. S., Braun, C., Birbaumer, N., Robinson, S. E., & Soekadar, S. R. (2016). Mapping entrained brain oscillations during transcranial alternating current stimulation (tACS). Neuroimage, 140, 89–98.

Zaehle, T., Rach, S., & Herrmann, C. S. (2010). Transcranial alternating current stimulation enhances individual alpha activity in human EEG. PloS one, 5(11), e13766.

